# Host Cell Rap1b mediates cAMP-dependent invasion by *Trypanosoma cruzi*

**DOI:** 10.1101/2022.06.22.497233

**Authors:** Gabriel Ferri, Martin M. Edreira

**Affiliations:** CONICET-Universidad de Buenos Aires, IQUIBICEN, Ciudad de Buenos Aires, Argentina; Laboratorio de Biología Molecular de Trypanosomas, Departamento de Química Biológica, Facultad de Ciencias Exactas y Naturales, Universidad de Buenos, Ciudad de Buenos Aires, Argentina

**Author notes:** **Correspondence:** Martin M. Edreira.

**Keywords:** Trypanosoma cruzi, cAMP signaling, Epac, PKA, Rap1b, MAK/ERK, Host cell invasion

## Abstract

*Trypanosoma cruzi* cAMP-mediated invasion has been long described, however, the detailed mechanism of action of the pathway activated by this cyclic nucleotide still remains unknown. We have recently demonstrated a crucial role for Epac in the cAMP-mediated invasion of the host cell. In this work, we proved that the cAMP/Epac pathway is activated in different cells lines and, by pull-down experiments designed to identify only the active form of Rap1b (Rap1b-GTP) and invasion assays using cells transfected with a constitutively active form of Rap1b (Rap1b-G12V), established the participation of Rap1b as mediator of the pathway. In addition to the activation of this small GTPase, fluorescence microscopy allowed us to demonstrate the relocalization of Rap1b to the entry site of the parasite. Moreover, phospho-mimetic and non-phosphorylable mutants of Rap1b were used to demonstrate a PKA-dependent antagonistic effect on the pathway, by phosphorylation of Rap1b, and potentially of Epac. Finally, Western Blot analysis was used to determine the involvement of the MEK/ERK signalling downstream of cAMP/Epac/Rap1b-mediated invasion.

## Introduction

As an obligate intracellular parasite, *Trypanosoma cruzi* replicates in the cytoplasm of infected mammalian host cells. Attachment of trypomastigotes activates several host signalling pathways, including the elevation of intracellular cAMP levels in the host cell (Ferri and Edreira, 2021). It has been shown that invasion involves the recruitment and fusion of lysosomes to the entry site (Andrews, 1995), and that cAMP potentiates the Ca^2+^-dependent exocytosis of lysosomes and lysosome-mediated cell invasion (Rodríguez et al., 1996). Although a transient increase of Ca^2+^ and the recruitment of lysosomes are common features in the invasion of metacyclic trypomastigotes (MTs) and tissue culture-derived trypomastigotes (TCTs) (Rodriguez et al., 1999; Martins et al., 2011), the signalling pathways that these parasites promote in the host cell are different. Among them, the activation of cAMP-mediated signalling by TCTs is a poorly studied process. It has been previously demonstrated that the pharmacologic intervention of the cAMP pathway was able to modulate parasite invasion (Rodriguez et al., 1999; Fernandes et al., 2006; Musikant et al., 2017). To determine the specific role of cAMP main effectors, PKA and Epac, in *T. cruzi* invasion, we used a set of pharmacological tools to selectively activate or inhibit these proteins. Whereas differential activation of PKA had no effect, a significant increase in invasion was observed in cells treated with a cAMP analogue that exclusively activates Epac (Musikant et al., 2017). Accordingly, inhibition of Epac by ESI-09 (Enserink et al., 2002) showed a significant decrease in invasion. Unexpectedly, specific inhibition of PKA also showed a positive effect on invasion, suggesting a PKA/Epac crosstalk during the process of invasion (Musikant et al., 2017). In this regard, it has been described that both proteins can be recruited to the same microdomain through the association with radixin (Gloerich et al., 2010; Hochbaum et al., 2011), an ERM structural protein that attaches the plasma membrane to the cortical actin cytoskeleton (McClatchey, 2014). Respectively, confocal studies have shown that ERM proteins are associated with the invasion site of extracellular amastigotes (EAs), where colocalize with F-actin (Ferreira et al., 2017). Moreover, a link between radixin and the cAMP/Epac-dependent pathway during TCT invasion was confirmed by blocking host cell invasion with a permeable version of 15-mer sequence (stearate-KPRACSYDLLLEHQRP-amide peptide) corresponding to the minimal Epac1 ERM binding domain. This peptide displaces the Epac protein from its association with radixin and delocalized it from the microdomain. Under these conditions, the percentage of invasion is similar to that obtained when the Epac protein is inhibited by ESI-09 (Musikant et al., 2017). Taken together, these results clearly established a crucial role for Epac in the cAMP-mediated invasion of the host cell. However, downstream effectors involved in this pathway are still unknown. Epac1 has been involved in PI3K/Akt and MEK/ERK pathways (Baviera et al., 2010; Gündüz et al., 2019), and members of these pathways, including Rap1, were localized at late endosomes/lysosomes (Pizon et al., 1994). In cardiomyocytes, the cAMP/Epac/Rap1 pathway modulates the excitation-contraction mechanism by stimulating Ca^2+^ release through ryanodine receptors (RyR) (Oestreich et al., 2009). In smooth muscle cells, Rap1 inhibits RhoA activity and promotes Ca^2+^ desensitization and smooth muscle relaxation (Zieba et al., 2011). Rap1 activation also induces muscle hyperpolarization by decreasing Ca^2+^ influx by inducing the opening of Ca^2+^-sensitive K^+^ channels, generating a boost in vasodilation (Kosuru and Chrzanowska, 2020). In addition, Rap1 was shown to modulate mitogen-activated kinases (MAPKs), like extracellular signal-regulated kinase (ERK1/2), inducing the stimulation or inhibition of these kinases depending on the cell type. More recently, the role of Rap1 in ERK phosphorylation and activation in smooth muscle was demonstrated (Li et al., 2018). Ral-GDS, an effector of Rap1, promotes cardiomyocyte autophagy (Rifki et al., 2013), and the downstream effector of Ral-GDS, RalB, binds specific subunits of the exocytosis machinery and mediates activation of autophagosome assembly (Bodemann et al., 2011).

Smooth muscle and heart are the most important target organs for *T. cruzi* infection and persistence during the chronic phase of Chagas disease. Taking into account that Epac has a critical role in cAMP-mediated invasion and the regulation of various cAMP-dependent functions in smooth muscle and heart, possibly modulating the intracellular concentration of Ca^2+^ through the activation of Rap1 and the participation of ERK1/2 (Ruiz-Hurtado et al., 2013; Lezoualc’H et al., 2016; Kosuru and Chrzanowska, 2020), deciphering the detailed functioning of the cAMP/Epac pathway would provide a deeper insight into the host cell invasion mechanisms mediated by this cyclic nucleotide. In this work, we investigated the involvement of two known effectors, Rap1b and ERK, as potential mediators in the cAMP/Epac-dependent invasion by *T. cruzi* and the role of PKA-dependent Rap1b phosphorylation.

## Materials and Methods

### Cells and parasites

NRK (ATCC® CRL-6509™), VERO (ATCC® CCL-81™) and HELA (ATCC® CCL-2™) cell lines were cultured in DMEM medium supplemented with Glutamax™ (Gibco), 10% (v/v) FBS (Natocor), 100 U/ml penicillin and 0.1 mg/ml streptomycin (Sigma), and maintained at 37°C in a 5% CO_2_ atmosphere. The HL-1 cell line (Claycomb et al., 1998) was cultured in a gelatin/fibronectin matrix (5 μg fibronectin / 0.02% gelatin (m/v)-Sigma) and Claycomb culture medium supplemented with Glutamax™ (Gibco), 10% (v/v) FBS, 100 U/ml penicillin, 0.1 mg/ml streptomycin and 0.1 mM norepinephrine (Sigma). Tissue culture-derived trypomastigotes forms (TCT) of *T. cruzi* Y strain were routinely maintained in VERO cells cultured in DMEM supplemented with 4% FBS and penicillin/streptomycin. Trypomastigotes were obtained from supernatants of infected VERO cells by centrifugation.

### Invasion assay

Cells were grown on glass cover slides with DMEM 10% FBS for 24 hours at 2×10^4^ cells/well density at 37°C, 5% CO^2^ and incubated with or without: 37,5 µM of the Epac1 inhibitor ESI-09 (Sigma); 300 µM of 8-Br-cAMP (Biolog); 50 µM of the MEK1/2 inhbitor PD98059. Cells were then washed and infected with trypomastigotes of the Y strain (moi 100:1) for 2 hours. Parasite were removed and cells incubated for 48 hs. Cells were fixed, stained with DAPI and infection level determined by fluorescence microscopy. Percentage of invasion and amastigotes/100 cells were calculated counting 3,000 cells. Infection of non-treated cells was considered as basal infection.

### Host cell transfection

A transient transfection protocol with polyethyleneimine (PEI) was used (Longo et al., 2013). Briefly, cells were grown at about 60% confluence and incubated at 37°C in a 5% CO_2_, 95% humidified air environment. Next day, cells were transfected with pCGN empty vector (EMPTY), pCGN-HA-Rap1b (HA-Rap1) or HA-Rap1b mutants (G12V, S179A, S179D, or combinations) (kindly provided by Dr D. Altschuler, University of Pittsburgh, USA) using a ratio of 4:1 DNA:PEI mix in OptiMEM medium (Gibco). The mixture was kept for 30 min. at room temperature and then added to the cells and incubated at 37°C and 5% CO_2_. After 24 h, cells were washed with PBS and complete medium (DMEM or Claycomb 10% FBS) was added. The transfected cells were used at 24h post-transfecction.

### Trypomastigote release assay

HL-1 cells were seeded on a 24-well plate at a concentration of 7000 cells/mL in Claycomb medium supplemented with 10% FBS. After 24 hours, cells were infected and treated as described above. 72 hours later, medium was replaced with fresh prepared treatments until trypomastigotes were observed under microscope at six days post infection (pi). Supernatants were transferred to a new plate and a solution of resazurin sodium salt dye was added (final concentration 0.1 mM). After 3 hours of incubation, fluorescence was measured with a FLUOstar OPTIMA (BMG LABTECH) microplate reader at 590 nm (excitation: 570 nm). Baseline corrected values of fluorescence were normalized to the negative control.

### GST Pull-down

A total of 1 mL bacteria lysates containing GST or GST-RBD were mixed by rotation with 40 µl 50% GSH-Sepharose at 4 °C for 1 h. The beads were centrifuged and washed with lysis buffer. Lysates from HA-Rap1 transfected cells pre-treated for 2h with 8Br-cAMP, infected with trypomastigotes of the Y strain (Tp Y) or mock infected (Ctrol) were incubated with RBD-glutathione-agarose resin for 1h at 4°C. Resin was washed and eluted with cracking buffer for WB analysis.

### Western Blot

After electrophoresis, the gel was equilibrated in 25 mM Trizma base, 192 mM L-1 glycine and 20% v/v methanol pH 8.3. Then, proteins were transferred to previously hydrated with methanol PVDF membranes (Amersham™ Hybond, GE Healthcare) in a vertical tank (Mini-PROTEAN® Tetra Cell, Bio-Rad). After transfer, membranes were blocked with 20 mM L−1 Tris-HCl, 500 mM NaCl, 0.05% Tween and 5% non-fat milk, pH 7.5, incubated with anti-GST (Genscript), anti-p44/42 MAPK (ERK1/2, Cell Signaling), anti-phospho-p44/42 MAPK (pERK1/2, Cell Signaling) or anti-GAPDH (Santa Cruz Biotechnology) antibodies. After incubation, membrane was washed and incubated with rabbit horseradish peroxidase (HRP)-IgGs antibody (Santa Cruz Biotechnology), washed again and then revealed using 0.88 mg/ml luminol, 0.066 mg/ml p-coumaric acid, 6 mM H_2_O_2_; 100 mM Tris-HCl, pH 8.8 solution. Chemiluminescence was recorded with the C-DiGit scanner (LI-COR), and bands intensity were quantified with ImageJ software.

### ERK phosphorylation

Cells were treated with or without PD98059 for 2h and incubated with trypomastigotes of the Y strain, treated with 750 µM H_2_O_2_ for 5 min or mock infected. Then, cells were lysed and cracking buffer added for WB analysis.

### Indirect immunofluorescence assay

Cells were adhered to glass previously treated with 40 μg/ml of poly-D-lysine (Sigma), fixed with PBS-PFA 4% (Sigma), washed with PBS and incubated with NH_4_Cl for 15 min. Then, were permeabilized with 0.2% Triton-x100 and incubated with anti-RAP1 antibody (Genscript) at 4°C. After 16 h, washed with PBS, incubated with mouse anti-IgG (H+L) anti-conjugated to Alexa Fluor®594 antibody (Jackson InmunoResearch), and nuclei stained with DAPI. Finally, glasses were mounted on slides with FluorSave™ mounting solution (Merk Millipore). Preparations were analysed in a Nikon Eclipse E600 fluorescence microscope.

## Results

### cAMP/Epac activation as a ubiquitous mechanism of invasion in *T. cruzi*

The crucial role of Epac during invasion by *T. cruzi* was recently described in NRK cells (Musikant et al., 2017). In order to assess the ubiquity of the cAMP/Epac pathway, other cell lines were used in invasion assays. Similar to what happened in NRK cells (Rodriguez et al., 1999; Musikant et al., 2017), high levels of cAMP induced by a non-hydrolysable permeable analogue of cAMP, 8-Br-cAMP (8-Bromoadenosine 3′,5′-cyclic monophosphate) (Biolog), positively modulated invasion in both HELA and HL-1 cells (Figure 1). Consistent with this result, specific pharmacological inhibition of Epac by ESI-09 (Sigma), resulted in a significant decrease in invasion in both cell lines (Figure 1).

**Figure 1:**
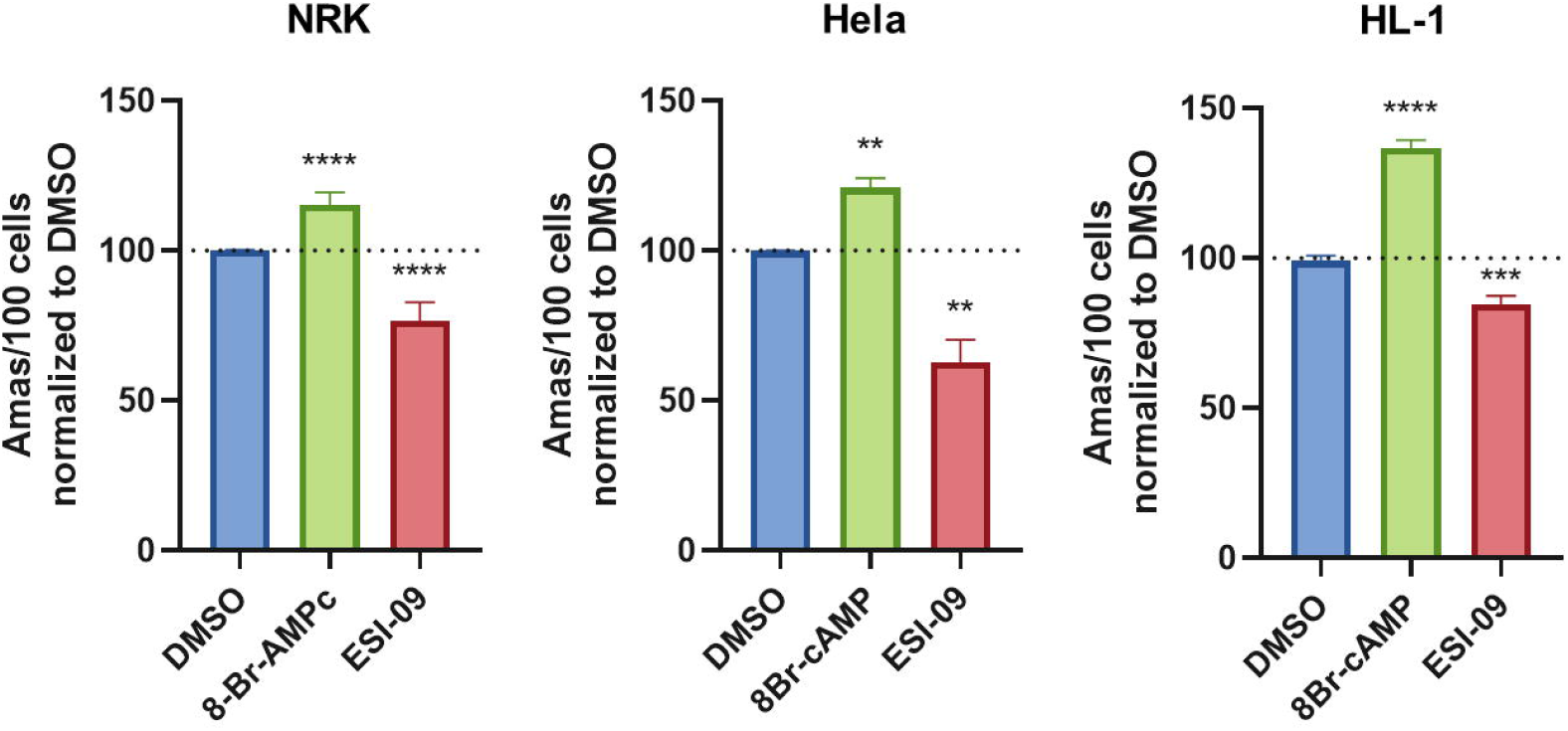
cAMP/Epac pathway is required for *T. cruzi* invasion in different cell lines. Pre-treated NRK, HELA or HL-1 cells (30 min at 300 μM of 8-Br-cAMP or 37.5 μM of ESI-09) were infected with trypomastigotes from *T. cruzi* Y strain (100:1 parasite to cell ratio for 2 h). 48 hs post-infection cells were fixed, stained with DAPI and percentage of invasion determined by fluorescence microscopy. Infection of untreated cells was considered as basal infection. Results are expressed as mean ± SD (n ≥ 3). **** p<0.0001, *** p <0.001, ** p <0.01; ANOVA and Dunnett’s post-test.

### Rap1b as a mediator of the cAMP/Epac-dependent invasion

The participation of Rap1b as mediator of the cAMP/Epac-dependent invasion was evaluated by pull-down experiments using agarose bound GST-RalGDS Rap-binding domain (GST-RBD) designed to pull down only the active form of Rap1b (Rap1b-GTP). Briefly, cells transfected with HA-Rap1b were incubated in the presence of DMSO (vehicle), 8-Br-cAMP or trypomastigotes of Y strain of *T. cruzi* for 2 h. Cells were lysed and lysates used in pull-down experiments. As shown in Figure 2, higher levels of activated GTP-bound Rap1 were detected in lysates from cells incubated with 8-Br-cAMP and trypomastigotes, supporting the involvement of Rap1b in cAMP-mediated invasion. In accordance with these results, HELA and HL-1 cells transfected with a constitutively active form of Rap1b, Rap1b-G12V, presented a significant increase in invasion, when compared with the control (Figure 3). Noteworthy, when the invasion-differentiation-release cycle was evaluated in HL-1 cells overexpressing Rap1b-G12V, trypomastigotes released into the medium reflected the results obtained for invasion, suggesting the cAMP/Epac/Rap1b pathway played a role in the early steps of the establishment of infection (Figure 3Biii). In addition to these results that demonstrated that Rap1b-GTP is required as a mediator of the cAMP/Epac1 pathway during the invasion by *T. cruzi*, fluorescence microscopy revealed the relocalization of Rap1b, reflected as an increase in the fluorescence intensity of Rap1, to the site of entry of *T. cruzi* (Figure 4), supporting the hypothesis that Rap1b needs to be activated and properly localized in the entry site.

**Figure 2:**
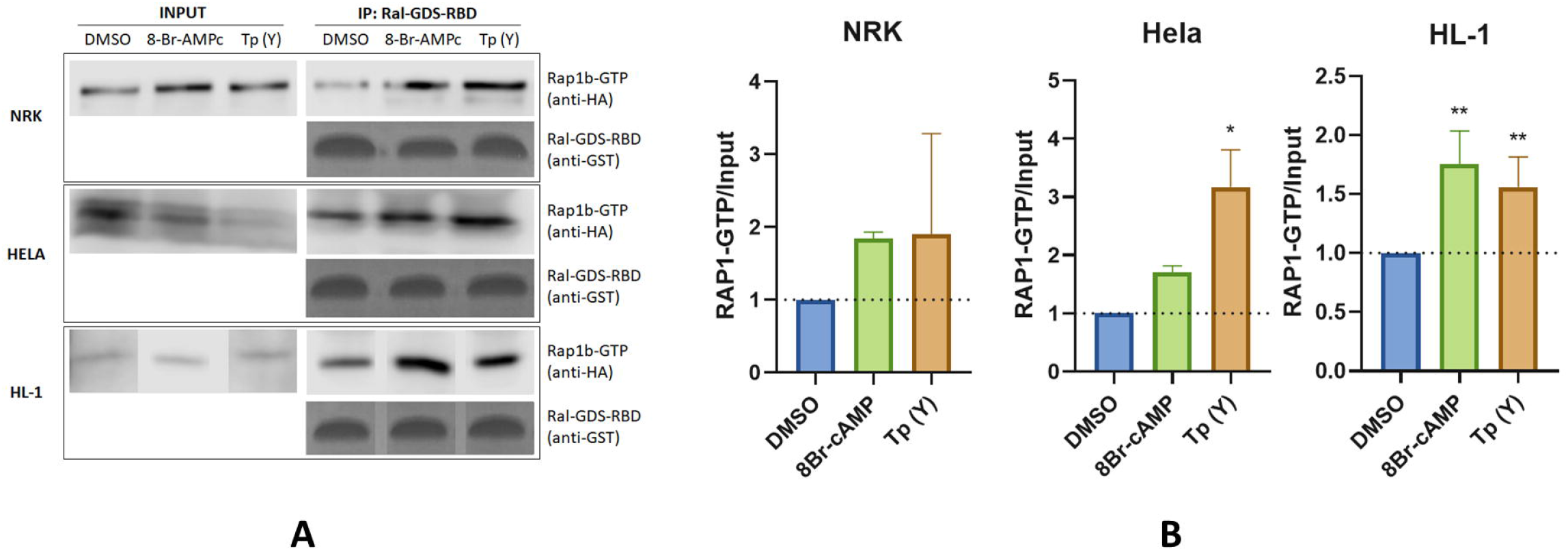
Rap1b pull-down assays. **A)** HA-Rap1 transfected NRK, HELA or HL-1 cells were incubated for 2 h with 8-Br-cAMP, infected with trypomastigotes from *T. cruzi* Y strain or mock infected (DMSO). Then, cells were lysed and pull-down assay with glutathione-agarose resin performed for 1 h at 4°C. Resin was washed and eluted with cracking buffer for WB analysis. **B)** Bands were quantified and normalized against the input using ImageJ cell software. Results are expressed as mean ± SD (n≥3). * p<0.05, ** p <0.005, One-way ANOVA – Dunnett’s multiple comparison test.

**Figure 3:**
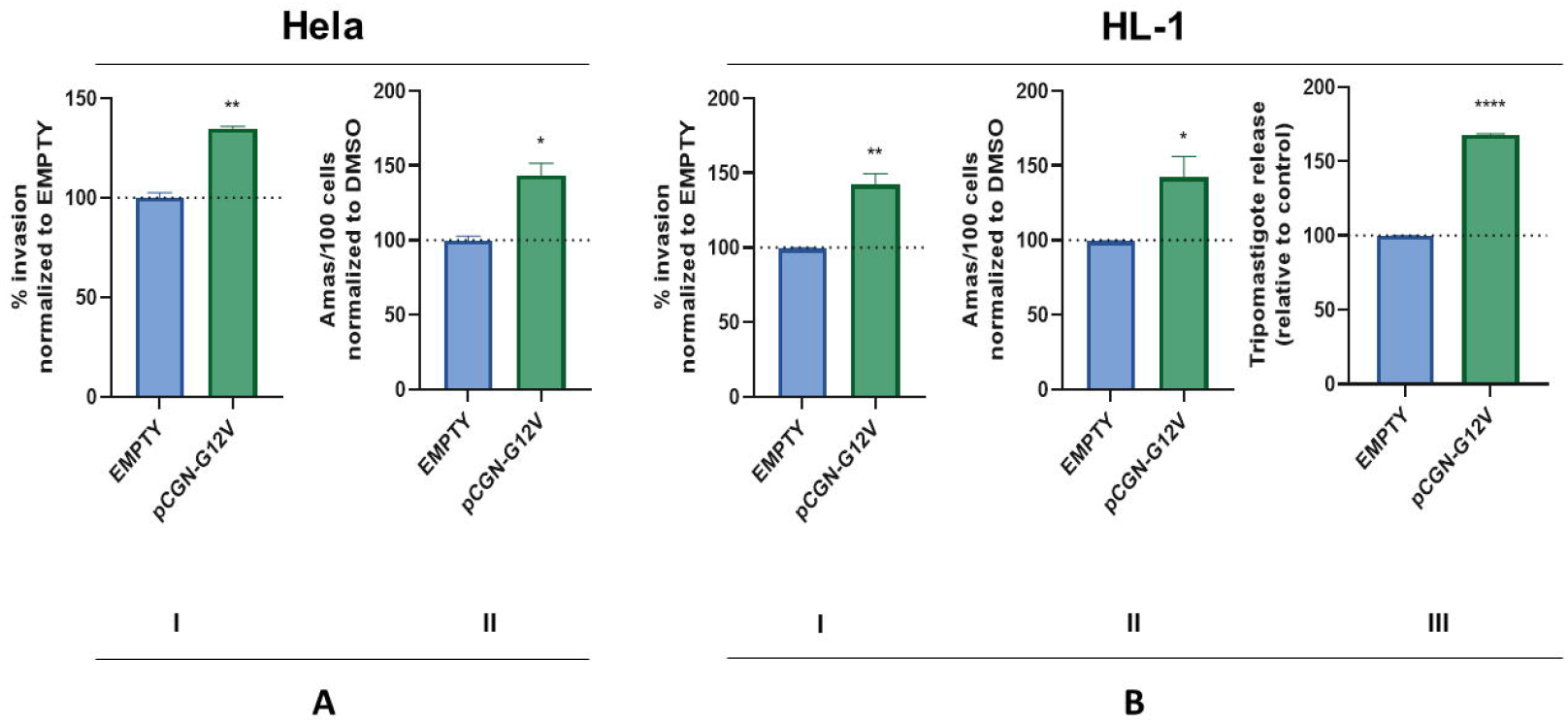
Invasion assays. Rap1 transfected Hela and HL-1 cells were infected with trypomastigotes from *T. cruzi* Y strain (100:1 parasite to cell ratio for 2 h). 48 hs post-infection cells were fixed, stained with DAPI and percentage of invasion determined by fluorescence microscopy. Infection of untreated cells was considered as basal infection. Results are expressed as mean ± SD (n ≥ 3), ** p <0.01, * p <0.1; ANOVA and Dunnett’s post-test. In the case of the trypomastigote release assay, HL-1 cells were infected and treated as described above. 72 hours later, medium was replaced with fresh prepared treatments until trypomastigotes were observed under microscope at six days post infection (pi). Supernatants were transferred to a new plate and quantification of trypomastigotes was performed with resazurin method. Results are expressed as mean ± SD (n ≥ 3). **** p<0.0001, t student test.

**Figure 4:**
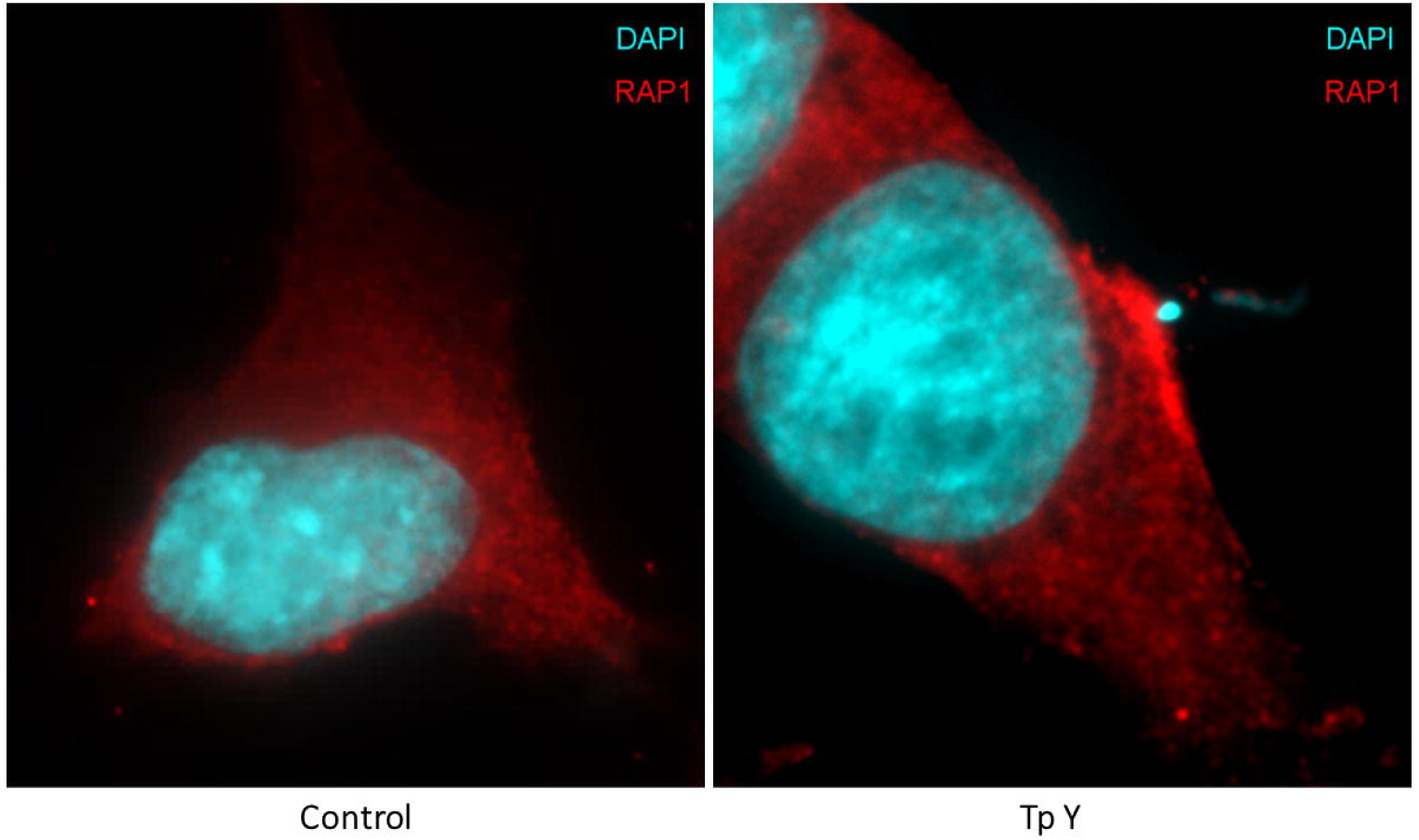
Immunofluorescence of HL-1 cells were infected for 5 to 15 min with (Tp Y) trypomastigotes from *T. cruzi* Y strain or mock infected (Control) and then fixed and incubated with primary antibody against Rap1 protein and a secondary antibody conjugated to Alexa594. Photos were taken with a fluorescence microscope.

### Role of PKA-dependent Rap1b phosphorylation

While the specific activation of PKA had no effect on invasion, an increase in invasion was observed as a result of PKA inhibition (Musikant et al., 2017). Therefore, under physiological conditions, PKA-mediated phosphorylation would negatively regulate the cAMP/Epac pathway of invasion. The inhibition of the Epac-mediated invasion pathway could be achieved, at least, at two different levels: through direct phosphorylation of Epac or at the level of Rap1, an Epac effector and a known target for PKA-mediated phosphorylation (Edreira et al., 2009). To evaluate the role of Rap1b phosphorylation, HELA cells overexpressing phospho-mimetic (S179D) or phospho-deficient (S179A) Rap1b mutants were infected and trypomastigotes released at day 6 pi were counted using resazurin method (Rolón et al., 2006). As shown in Figure 5, cells transfected with the phospho-mimetic Rap1b-S179D mutant presented a decreased invasion with respect to control cells or cells overexpressing the non-phosphorylable mutant Rap1b-S179A, supporting a PKA-dependent antagonistic effect on the pathway. Interestingly, the effect of phosphorylation could be reverted by transfecting cells with the double mutant G12V/S179D, a constitutive active phospho-mimetic Rap1, opening the possibility of a two-level regulation of PKA on the Epac/Rap1 pathway.

**Figure 5:**
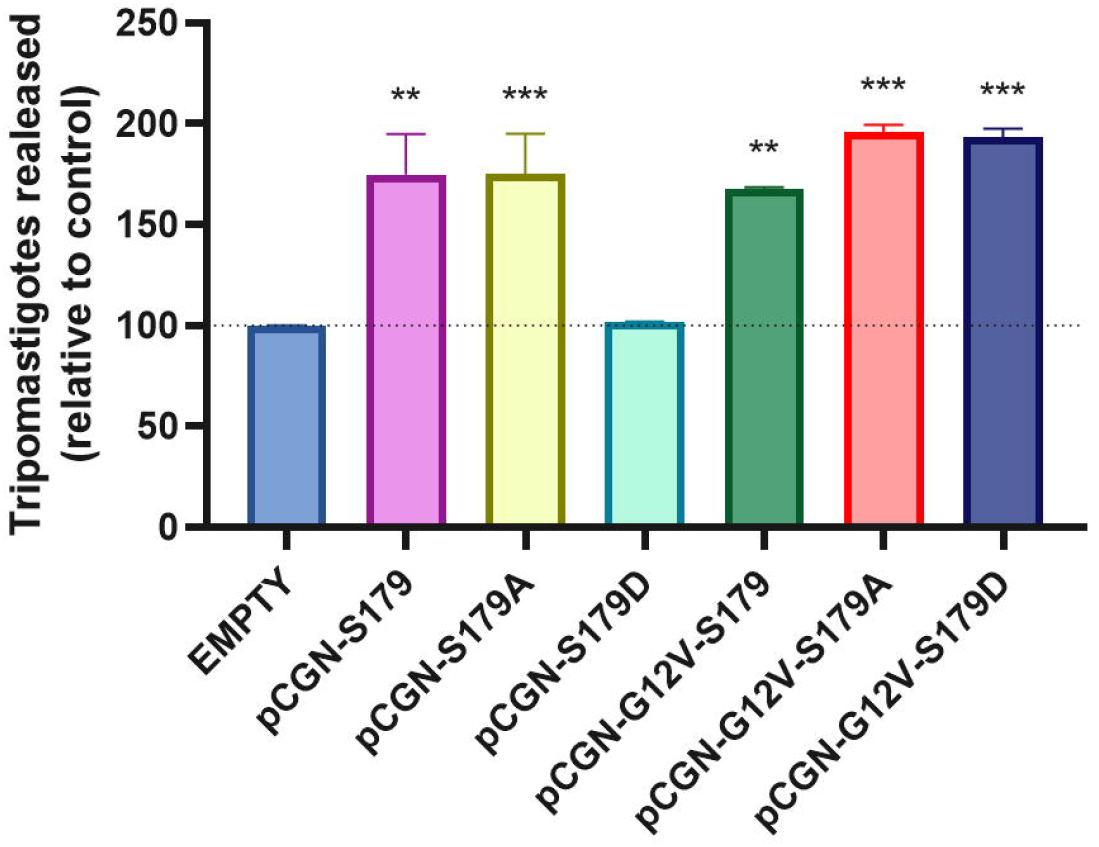
PKA-dependent phosphorylation effect on invasion. Transfected HL-1 cells were infected and treated as described above. 72 hours later, medium was replaced with fresh prepared treatments until trypomastigotes were observed under microscope at six days post infection (pi). Supernatants were transferred to a new plate and quantification of trypomastigotes was performed with resazurin method. Results are expressed as mean ± SD (n ≥ 3), *** p <0.001, ** p <0.01; ANOVA and Dunnett’s post-test.

### MEK/ERK as a downstream effector of cAMP/Epac-mediated invasion of *T. cruzi*

In order to elucidate the involvement of MEK/ERK in the cAMP-dependent invasion, activation of ERK1/2 was analysed by Western Blot. An increase in ERK1/2 phosphorylation in both NRK and HL-1 cells was observed during the host cell invasion (Figure 6A). To determine whether the activation of ERK1/2 modulates the invasion levels, cells pre-treated with the MEK1/2 kinase inhibitor PD98059 were infected with the parasite. In accordance, the inhibition of ERK phosphorylation produced a significant decrease in invasion (Figure 6B). MEK/ERK could be independently activated or a downstream effector of Epac/Rap1. The fact that the inhibition of MEK or Epac induced a similar decrease in invasion, and no additive or synergic effects were observed when both proteins were simultaneously inhibited (Figure 6C), suggests that MEK/ERK is a downstream effector of cAMP/Epac1/Rap1b-mediated invasion.

**Figure 6:**
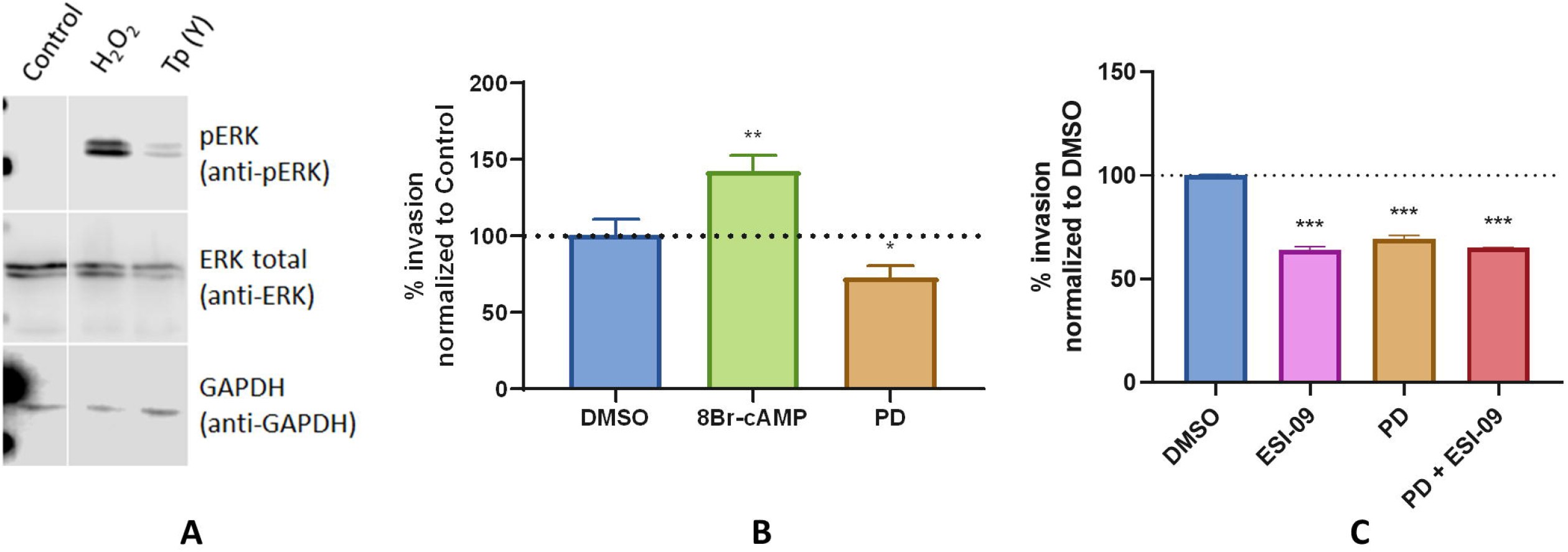
ERK phosphorylation. **A)** NRK or HL-1 cells were incubated for 2 h with trypomastigotes from *T. cruzi* Y strain (Tp Y), treated with 750 μM H_2_O_2_ for 5 min (positive control) or mock infected (Control). Then, cells were lysed and cracking buffer added for WB analysis. **B)** and **C)** Invasion assay. Pretreated HELA cells were infected with trypomastigotes from *T. cruzi* Y strain (100:1 parasite to cell ratio for 2 h). 48 hs post-infection cells were fixed, stained with DAPI and percentage of invasion determined by fluorescence microscopy. Infection of untreated cells was considered as basal infection. Results are expressed as mean ± SD (n ≥ 3) *** p <0.001, ** p <0.01; ANOVA and Dunnett’s post-test.

## Discussion

*T. cruzi* invasion showed to be a complex process just taking into account the different stages of the parasite with the ability to infect host cells. This complexity is even higher when considering different DTUs, strains, the repertoire of surface/secreted molecules and the signalling pathways activated in the host cell (Ferri and Edreira, 2021). Despite being able to infect any nucleated cell, it has been shown that *T. cruzi* exhibits a certain cellular tropism (Santi-Rocca et al., 2017) and that the signalling pathways activated in the host cell differ according to the stage of the parasite (Maeda et al., 2012). In this context, it has been reported that the activation of cAMP-mediated signalling pathways triggers Ca^2+^-dependent lysosomal exocytosis and promotes host cell invasion by *T. cruzi* (Rodriguez et al., 1999). The Ca^2+^ release from intracellular compartments, such as the endoplasmic reticulum, is associated with an increase in intracellular levels of cAMP. In mammalian cells, cAMP downstream effectors, PKA and Epac, are involved in Ca^2+^-activated exocytosis events (Seino and Shibasaki, 2005). Furthermore, members of these pathways, including Rap1, have been localized to late endosomes/lysosomes (Pizon et al., 1994), and Epac-mediated activation of Rap1 has been identified in regulated exocytosis in human sperm (Miro-Moran et al., 2012), insulin secretion (Tengholm and Gylfe, 2017), and pancreatic amylase release (Sabbatini et al., 2008). It was previously shown that Epac1-mediated signalling represents the main mechanism for cAMP-mediated invasion by *T. cruzi* (Musikant et al., 2017). In addition, ERM proteins, which are essential for the function and architecture of the cell cortex by linking the plasma membrane to the underlying actin cytoskeleton (McClatchey, 2014), have been associated with the invasion of EAs (Ferreira et al., 2017). Moreover, in confocal studies, it was shown that ERM proteins are recruited at the entry site of the parasites where they colocalize with F-actin, while its depletion inhibits HELA cells invasion (Ferreira et al., 2017). Remarkably, one of its members, radixin, was identified as a scaffold unit for cAMP effectors in the spatial regulation of Epac1/Rap1-mediated signaling (Gloerich et al., 2010; Hochbaum et al., 2011). In this regard, we have previously revealed a link between Epac1 and radixin in the cAMP-mediated invasion of TCTs, by blocking the invasion of NRK cells with a permeable peptide of 15 amino acids that binds to the minimal ERM-binding domain of Epac (Musikant et al., 2017). In order to elucidate the role of cAMP downstream effectors involved in *T. cruzi* invasion, we evaluated the activation of the cAMP/Epac pathway by TCTs of Y strain in NRK, HELA and HL-1 cell lines. NRK cells are normal fibroblasts from rat kidney, originally used in the establishment of cAMP as a modulator of invasion events (Rodriguez et al., 1999) and to demonstrate the participation of Epac1 as the main effector of this modulation (Musikant et al., 2017). On the other hand, HELA cells are epithelial human cervix cells that have been widely used in invasion assays (Clemente et al., 2016; Ferreira et al., 2019; Rodrigues et al., 2019) and HL-1 cells, previously used in invasion assays, as well (Benatar et al., 2015), are cardiomyocytes from mouse heart, one of the most important target organs in the infection and persistence of *T. cruzi*. The data obtained showed that the activation of the cAMP/Epac pathway by TCTs occurs regardless of the origin (rat, mouse, human) or the cell type (kidney, cervix, heart) that the parasite is invading. In addition, we investigated the role of Rap1b during the cAMP/Epac1-mediated invasion. Rap1b, a GTPase of the Ras family, is known to integrate Epac-and/or PKA-dependent events to achieve an efficient cAMP signal transduction (Hochbaum et al., 2008; Jaśkiewicz et al., 2018; Kosuru and Chrzanowska, 2020). Pull-down assays were used to detect higher levels of activated GTP-bound Rap1 in lysates from infected cells. Likewise, cells transfected with the constitutively active form of Rap1b (G12V) were more susceptible to invasion, compared to the control. Moreover, from fluorescence microscopy assays it was evident the recruitment of Rap1b to the parasite entry site. Interestingly, when studying PKA participation using a specific inhibitor of this kinase, it was observed that the invasion levels of TCTs increased compared to the control (Musikant et al., 2017), suggesting a PKA-dependent antagonist effect. This effect could be mediated by PKA phosphorylation of the effectors of the cAMP pathway, such as Epac and Rap1b. PKA-dependent phosphorylation at S179 of Rap1b has long been established (Altschuler and Lapetina, 1993). Our results support the antagonistic effect of PKA through, at least, Rap1b phosphorylation, since invasion was affected in cells transfected with phospho-mimetic Rap1b-S179D, with respect to control cells and cells overexpressing Rap1b-S179A, the non-phosphorylable version of Rap1b. In line with these observations, it has been shown that Rap1b phosphorylation destabilizes the association of this protein with the plasma membrane and promotes Rap1b inactivation (Ntantie et al., 2013; Takahashi et al., 2013). Here, we demonstrated that the activation of Rap1b is required during the TCT invasion as a mediator of the cAMP/Epac1 pathway and that Rap1b relocalized to the entry site of the parasite. However, PKA negative effect on invasion was abrogated in the presence of the constitutively active G12V mutation, suggesting that Rap1b is required in the phosphorylated and inactive form to completely abolish the cAMP/Epac/Rap1b pathway and, thus, potential regulation of PKA over Epac.

It has been described that the MEK/ERK pathway can be activated or inhibited by cAMP (Stork and Schmitt, 2002). Furthermore, the activation of this pathway participates in the invasion of *T. cruzi* by way of the interaction of the host cell with parasite surface molecules, such as TS (Chuenkova and Pereira, 2001), Tc85 (Magdesian et al., 2007) or TSSAII (Cánepa et al., 2012). Also, Rap1 is associated with the phosphorylation and activation of ERK1/2 in smooth muscle (Li et al., 2018). Accordingly, our data revealed that TCTs induce ERK1/2 phosphorylation in mammalian cells and ERK1/2 activation modulates the invasion of these parasites as a downstream effector of Epac/Rap1-mediated invasion.

Although the transient increase in cytosolic Ca^2+^ concentration and lysosome recruitment that occur during invasion are characteristics shared between MTs and TCTs (Rodriguez et al., 1999; Martins et al., 2011), the signalling pathways triggered by both forms of parasites in the host cell are different. In TCT invasion, ERK1/2 activation is a distinctive feature that is mediated by Ca^2+^-dependent lysosomal exocytosis through the regulation of F-actin and the activation of the focal adhesion kinase (FAK) (Onofre et al., 2019). During MT invasion, in contrast, PKC promotes Ca^2+^ release from inositol 3-phosphate (IP3)-sensitive compartments through the binding of the surface glycoprotein gp82 to LAMP-2 receptors (Maeda et al., 2012; Onofre et al., 2021). On the contrary, the activity of PKC is not require for the invasion of TCTs in NRK cells, since treatment with PKC inhibitors did not affect the response to Ca^2+^ or the reorganization of F-actin, and has no effect on parasite internalization (Rodriguez et al., 1995). The divergence between the signalling pathways triggered by MTs and TCTs might be associated with the fact that the internalization of TCT is initiated by an invagination of the plasma membrane (Woolsey et al., 2003), in a lysosomal exocytosis-dependent process induced by a membrane injury and the following activation of the PMR mechanism (Fernandes et al., 2011). These mechanisms lead to changes that take place through the inhibition of the Rho/Rho signaling pathway by PKA (Woolsey and Burleigh, 2004; Mott et al., 2009). The fact that RhoA promotes actin polymerization but has a negative effect on EAs internalization during HELA cell invasion (Bonfim-Melo et al., 2018) and that Rap1b inhibits RhoA/ROCK activity in the muscle smooth tissue (Lakshmikanthan et al., 2014), suggest the hypothesis that the cAMP/Epac1/Rap1b signalling pathway could be activated in the first steps of the invasion by *T. cruzi*, promoting Ca^2+^-dependent lysosomal exocytosis and the reorganization of the cytoskeleton. Once the parasite is inside the cell, a PKA-mediated inhibition of Epac/Rap1b might be necessary for the parasite retention. In accordance, our results showed that Rap1b seems to be associated with the plasma membrane at the parasite entry site where it could be required during the internalization process and PKA had an antagonistic effect, probably through the phosphorylation of the S179 of Rap1b.

In this work, in order to elucidate the mechanisms of cAMP-mediated invasion by *T. cruzi*, it has been shown that the Epac/Rap1b/ERK pathway is activated during host cell invasion and that it is negatively regulated by PKA, possibly through the phosphorylation of Epac and/or Rap1b. In addition, a detailed characterization of effectors involved in *T. cruzi* invasion would provide an attractive set of new therapeutic targets for the repositioning or the development of new antiparasitic drugs, since there is a large variety of therapies that target cAMP-mediated signalling (Parnell et al., 2015).

## Author Contributions

GF and MME conceived, planned, and designed experiments. GF conducted experiments. GF and MME analysed the data and wrote the manuscript. All authors contributed to the article and approved the submitted version.

## References

Altschuler, D., and Lapetina, E. G. (1993). Mutational analysis of the cAMP-dependent protein kinase-mediated phosphorylation site of Rap1b. J. Biol. Chem. 268, 7527–7531. doi: 10.1016/s0021-9258(18)53207-5.

Andrews, N. W. (1995). Lysosome recruitment during host cell invasion by Trypanosoma cruzi. Trends Cell Biol 5, 133–137. doi: S0962892400889655 [pii].

Baviera, A. M., Zanon, N. M., Navegantes, L. C. C., and Kettelhut, I. C. (2010). Involvement of cAMP/Epac/PI3K-dependent pathway in the antiproteolytic effect of epinephrine on rat skeletal muscle. Mol. Cell. Endocrinol. 315, 104–112. doi: 10.1016/j.mce.2009.09.028.

Benatar, A. F., García, G. A., Bua, J., Cerliani, J. P., Postan, M., Tasso, L. M., et al. (2015). Galectin-1 Prevents Infection and Damage Induced by Trypanosoma cruzi on Cardiac Cells. PLoS Negl. Trop. Dis. 9, 1–23. doi: 10.1371/journal.pntd.0004148.

Bodemann, B. O., Orvedahl, A., Cheng, T., Ram, R. R., Ou, Y. H., Formstecher, E., et al. (2011). RalB and the exocyst mediate the cellular starvation response by direct activation of autophagosome assembly. Cell. doi: 10.1016/j.cell.2010.12.018.

Bonfim-Melo, A., Ferreira, É. R., and Mortara, R. A. (2018). Rac1/WAVE2 and Cdc42/N-WASP participation in actin-dependent host cell invasion by extracellular amastigotes of Trypanosoma cruzi. Front. Microbiol. 9, 1–14. doi: 10.3389/fmicb.2018.00360.

Cánepa, G. E., Degese, M. S., Budu, A., Garcia, C. R. S., and Buscaglia, C. A. (2012). Involvement of TSSA (trypomastigote small surface antigen) in Trypanosoma cruzi invasion of mammalian cells 1. Biochem. J. 444, 211–218. doi: 10.1042/BJ20120074.

Chuenkova, M. V., and Pereira, M. A. (2001). The T. cruzi trans-sialidase induces PC12 cell differentiation via MAPK/ERK pathway. Neuroreport 12, 3715–3718. doi: 10.1097/00001756-200112040-00022.

Claycomb, W. C., Lanson, N. A. J., Stallworth, B. S., Egeland, D. B., Delcarpio, J. B., Bahinski, A., et al. (1998). HL-1 cells: a cardiac muscle cell line that contracts and retains phenotypic characteristics of the adult cardiomyocyte. Proc. Natl. Acad. Sci. U. S. A. 95, 2979–2984. doi: 10.1073/pnas.95.6.2979.

Clemente, T. M., Cortez, C., Novaes, A. da S., and Yoshida, N. (2016). Surface Molecules Released by Trypanosoma cruzi Metacyclic Forms Downregulate Host Cell Invasion. PLoS Negl. Trop. Dis. 10, 1–18. doi: 10.1371/journal.pntd.0004883.

Edreira, M. M., Li, S., Hochbaum, D., Wong, S., Gorfe, A. A., Ribeiro-Neto, F., et al. (2009). Phosphorylation-induced Conformational Changes in Rap1b. J. Biol. Chem. 284, 27480–27486. doi: 10.1074/jbc.M109.011312.

Enserink, J. M., Christensen, A. E., De Rooij, J., Van Triest, M., Schwede, F., Genieser, H. G., et al. (2002). A novel Epac-specific cAMP analogue demonstrates independent regulation of Rap1 and ERK. Nat. Cell Biol. 4, 1–6. doi: 10.1038/ncb874.

Fernandes, A. B., Neira, I., Ferreira, A. T., and Mortara, R. A. (2006). Cell invasion by Trypanosoma cruzi amastigotes of distinct infectivities: studies on signaling pathways. Parasitol Res 100, 59–68. doi: 10.1007/s00436-006-0236-6.

Fernandes, M., Cortez, M., Flannery, A. R., Tam, C., Mortara, R. A., and Andrews, N. W. (2011). Trypanosoma cruzi subverts the sphingomyelinase-mediated plasma membrane repair pathway for cell invasion. J. Exp. Med. 208, 909–921. doi: 10.1084/jem.20102518.

Ferreira, B. L., Ferreira, É. R., Bonfim-Melo, A., Mortara, R. A., and Bahia, D. (2019). Trypanosoma cruzi extracellular amastigotes selectively trigger the PI3K/Akt and Erk pathways during HeLa cell invasion. Microbes Infect. 21, 485–489. doi: 10.1016/j.micinf.2019.06.003.

Ferreira, É. R., Bonfim-Melo, A., Cordero, E. M., and Mortara, R. A. (2017). ERM proteins play distinct roles in cell invasion by extracellular amastigotes of Trypanosoma cruzi. Front. Microbiol. 8, 1–11. doi: 10.3389/fmicb.2017.02230.

Ferri, G., and Edreira, M. M. (2021). All Roads Lead to Cytosol: Trypanosoma cruzi Multi-Strategic Approach to Invasion. doi: 10.3389/fcimb.2021.634793.

Gloerich, M., Ponsioen, B., Vliem, M. J., Zhang, Z., Zhao, J., Kooistra, M. R., et al. (2010). Spatial Regulation of Cyclic AMP-Epac1 Signaling in Cell Adhesion by ERM Proteins. Mol. Cell. Biol. 30, 5421–5431. doi: 10.1128/mcb.00463-10.

Gündüz, D., Troidl, C., Tanislav, C., Rohrbach, S., Hamm, C., and Aslam, M. (2019). Role of PI3K/Akt and MEK/ERK Signalling in cAMP/Epac-Mediated Endothelial Barrier Stabilisation. Front. Physiol. 10, 1–13. doi: 10.3389/fphys.2019.01387.

Hochbaum, D., Barila, G., Ribeiro-Neto, F., and Altschuler, D. L. (2011). Radiin assembles cAMP effectors Epac and PKA into a functional cAMP compartment: role in cAMP-dependent cell proliferation. J Biol Chem 286, 859–866. doi: 10.1074/jbc.M110.163816.

Hochbaum, D., Hong, K., Barila, G., Ribeiro-Neto, F., and Altschuler, D. L. (2008). Epac, in synergy with cAMP-dependent protein kinase (PKA), is required for cAMP-mediated mitogenesis. J Biol Chem 283, 4464–4468. doi: C700171200 [pii] 10.1074/jbc.C700171200.

Jaśkiewicz, A., Pajką, B., and Orzechowski, A. (2018). The many faces of rap1 GTPase. Int. J. Mol. Sci. 19, 1–20. doi: 10.3390/ijms19102848.

Kosuru, R., and Chrzanowska, M. (2020). Integration of rap1 and calcium signaling. Int. J. Mol. Sci. 21, 1–21. doi: 10.3390/ijms21051616.

Lakshmikanthan, S., Zieba, B. J., Ge, Z.-D., Momotani, K., Zheng, X., Lund, H., et al. (2014). Rap1b in Smooth Muscle and Endothelium Is Required for Maintenance of Vascular Tone and Normal Blood Pressure. Arterioscler. Thromb. Vasc. Biol. 34, 1486–1494. doi: 10.1161/ATVBAHA.114.303678.

Lezoualc’H, F., Fazal, L., Laudette, M., and Conte, C. (2016). Cyclic AMP sensor EPAC proteins and their role in cardiovascular function and disease. Circ. Res. doi: 10.1161/CIRCRESAHA.115.306529.

Li, Q., Teng, Y., Wang, J., Yu, M., Li, Y., and Zheng, H. (2018). Rap1 promotes proliferation and migration of vascular smooth muscle cell via the ERK pathway. Pathol. - Res. Pract. 214, 1045–1050. doi: 10.1016/j.prp.2018.04.007.

Longo, P. A., Kavran, J. M., Kim, M. S., and Leahy, D. J. (2013). “Transient mammalian cell transfection with polyethylenimine (PEI),” in Methods in Enzymology (Academic Press Inc.), 227–240. doi: 10.1016/B978-0-12-418687-3.00018-5.

Maeda, F. Y., Cortez, C., and Yoshida, N. (2012). Cell signaling during Trypanosoma cruzi invasion. Front Immunol 3, 361. doi: 10.3389/fimmu.2012.00361.

Magdesian, M. H., Tonelli, R. R., Fessel, M. R., Silveira, M. S., Schumacher, R. I., Linden, R., et al. (2007). A conserved domain of the gp85/trans-sialidase family activates host cell extracellular signal-regulated kinase and facilitates Trypanosoma cruzi infection. Exp. Cell Res. 313, 210–218. doi: https://doi.org/10.1016/j.yexcr.2006.10.008.

Martins, R. M., Alves, R. M., Macedo, S., and Yoshida, N. (2011). Starvation and rapamycin differentially regulate host cell lysosome exocytosis and invasion by Trypanosoma cruzi metacyclic forms. Cell. Microbiol. 13, 943–954. doi: 10.1111/j.1462-5822.2011.01590.x.

McClatchey, A. I. (2014). ERM proteins at a glance. J. Cell Sci. 127, 3199–3204. doi: 10.1242/jcs.098343.

Miro-Moran, A., Jardin, I., Ortega-Ferrusola, C., Salido, G. M., Peña, F. J., Tapia, J. A., et al. (2012). Identification and function of exchange proteins activated directly by cyclic AMP (Epac) in mammalian spermatozoa. PLoS One. doi: 10.1371/journal.pone.0037713.

Mott, A., Lenormand, G., Costales, J., Fredberg, J. J., and Burleigh, B. A. (2009). Modulation of host cell mechanics by Trypanosoma cruzi. J Cell Physiol 218, 315–322. doi: 10.1002/jcp.21606.

Musikant, D., Ferri, G., Durante, I. M., Buscaglia, C. A., Altschuler, D. L., and Edreira, M. M. (2017). Host Epac1 is required for cAMP-mediated invasion by Trypanosoma cruzi. Mol. Biochem. Parasitol. 211, 67–70. doi: 10.1016/j.molbiopara.2016.10.003.

Ntantie, E., Gonyo, P., Lorimer, E. L., Hauser, A. D., Schuld, N., McAllister, D., et al. (2013). An Adenosine-Mediated Signaling Pathway Suppresses Prenylation of the GTPase Rap1B and Promotes Cell Scattering. Sci. Signal. 6. doi: 10.1126/scisignal.2003374.

Oestreich, E. A., Malik, S., Goonasekera, S. A., Blaxall, B. C., Kelley, G. G., Dirksen, R. T., et al. (2009). Epac and phospholipase Cε regulate Ca2+ release in the heart by activation of protein kinase Cε and calcium-calmodulin kinase II. J. Biol. Chem. 284, 1514–1522. doi: 10.1074/jbc.M806994200.

Onofre, T. S., Rodrigues, J. P. F., Shio, M. T., Macedo, S., Juliano, M. A., and Yoshida, N. (2021). Interaction of Trypanosoma cruzi Gp82 With Host Cell LAMP2 Induces Protein Kinase C Activation and Promotes Invasion. Front. Cell. Infect. Microbiol. 11, 1–16. doi: 10.3389/fcimb.2021.627888.

Onofre, T. S., Rodrigues, J. P. F., and Yoshida, N. (2019). Depletion of host cell focal adhesion kinase increases the susceptibility to invasion by trypanosoma cruzi metacyclic forms. Front. Cell. Infect. Microbiol. 9, 1–10. doi: 10.3389/fcimb.2019.00231.

Parnell, E., Palmer, T. M., and Yarwood, S. J. (2015). The future of EPAC-targeted therapies: agonism versus antagonism. Trends Pharmacol Sci 36, 203–214. doi: 10.1016/j.tips.2015.02.003.

Pizon, V., Desjardins, M., Bucci, C., Parton, R. G., and Zerial, M. (1994). Association of Rap1a and Rap1b proteins with late endocytic/phagocytic compartments and Rap2a with the Golgi complex. J Cell Sci 107 (Pt 6, 1661–1670. Available at: http://www.ncbi.nlm.nih.gov/entrez/query.fcgi?cmd=Retrieve&db=PubMed&dopt=Citation&list_uids=7962206.

Rifki, O. F., Bodemann, B. O., Battiprolu, P. K., White, M. A., and Hill, J. A. (2013). RalGDS-dependent cardiomyocyte autophagy is required for load-induced ventricular hypertrophy. J Mol Cell Cardiol 59, 128–138. doi: 10.1016/j.yjmcc.2013.02.015.

Rodrigues, J. P. F., Souza Onofre, T., Barbosa, B. C., Ferreira, É. R., Bonfim-Melo, A., and Yoshida, N. (2019). Host cell protein LAMP-2 is the receptor for Trypanosoma cruzi surface molecule gp82 that mediates invasion. Cell. Microbiol. 21, 1–11. doi: 10.1111/cmi.13003.

Rodriguez, A., Martinez, I., Chung, A., Berlot, C. H., and Andrews, N. W. (1999). cAMP regulates Ca2+-dependent exocytosis of lysosomes and lysosome-mediated cell invasion by trypanosomes. J Biol Chem 274, 16754–16759. Available at: http://www.ncbi.nlm.nih.gov/entrez/query.fcgi?cmd=Retrieve&db=PubMed&dopt=Citation&list_uids=10358016.

Rodriguez, A., Rioult, M. G., Ora, A., and Andrews, N. W. (1995). A trypanosome-soluble factor induces IP3 formation, intracellular Ca2+ mobilization and microfilament rearrangement in host cells. J. Cell Biol. 129, 1263–1273. doi: 10.1083/jcb.129.5.1263.

Rodríguez, A., Samoff, E., Rioult, M. G., Chung, A., and Andrews, N. W. (1996). Host cell invasion by trypanosomes requires lysosomes and microtubule/kinesin-mediated transport. J. Cell Biol. 134, 349–362. doi: 10.1083/jcb.134.2.349.

Rolón, M., Vega, C., Escario, J. A., and Gómez-Barrio, A. (2006). Development of resazurin microtiter assay for drug sensibility testing of Trypanosoma cruzi epimastigotes. Parasitol. Res. doi: 10.1007/s00436-006-0126-y.

Ruiz-Hurtado, G., Morel, E., Domínguez-Rodríguez, A., Llach, A., Lezoualc’h, F., Benitah, J. P., et al. (2013). Epac in cardiac calcium signaling. J. Mol. Cell. Cardiol. doi: 10.1016/j.yjmcc.2012.11.021.

Sabbatini, M. E., Chen, X., Ernst, S. A., and Williams, J. A. (2008). Rap1 activation plays a regulatory role in pancreatic amylase secretion. J Biol Chem 283, 23884–23894. doi: M800754200 [pii] 10.1074/jbc.M800754200.

Santi-Rocca, J., Fernandez-Cortes, F., Chillón-Marinas, C., González-Rubio, M. L., Martin, D., Gironès, N., et al. (2017). A multi-parametric analysis of Trypanosoma cruzi infection: Common pathophysiologic patterns beyond extreme heterogeneity of host responses. Sci. Rep. doi: 10.1038/s41598-017-08086-8.

Seino, S., and Shibasaki, T. (2005). PKA-dependent and PKA-independent pathways for cAMP-regulated exocytosis. Physiol Rev 85, 1303–1342. doi: 85/4/1303 [pii] 10.1152/physrev.00001.2005.

Stork, P. J.., and Schmitt, J. M. (2002). Crosstalk between cAMP and MAP kinase signaling in the regulation of cell proliferation. Trends Cell Biol 12, 258–266. doi: S0962892402022948 [pii].

Takahashi, M., Dillon, T. J., Liu, C., Kariya, Y., Wang, Z., and Stork, P. J. S. (2013). Protein kinase a-dependent phosphorylation of Rap1 regulates its membrane localization and cell migration. J. Biol. Chem. 288, 27712–27723. doi: 10.1074/jbc.M113.466904.

Tengholm, A., and Gylfe, E. (2017). cAMP signalling in insulin and glucagon secretion. Diabetes, Obes. Metab. doi: 10.1111/dom.12993.

Woolsey, A. M., and Burleigh, B. A. (2004). Host cell actin polymerization is required for cellular retention of Trypanosoma cruzi and early association with endosomal/lysosomal compartments. Cell Microbiol 6, 829–838. doi: 10.1111/j.1462-5822.2004.00405.x.

Woolsey, A. M., Sunwoo, L., Petersen, C. A., Brachmann, S. M., Cantley, L. C., and Burleigh, B. A. (2003). Novel Pl 3-kinase-dependent mechanisms of trypanosome invasion and vacuole maturation. J. Cell Sci. 116, 3611–3622. doi: 10.1242/jcs.00666.

Zieba, B. J., Artamonov, M. V, Jin, L., Momotani, K., Ho, R., Franke, A. S., et al. (2011). The cAMP-responsive Rap1 Guanine Nucleotide Exchange Factor, Epac, Induces Smooth Muscle Relaxation by Down-regulation of RhoA Activity. J. Biol. Chem. 286, 16681–16692. doi: 10.1074/jbc.M110.205062.

